# TRIM34 acts with TRIM5 to restrict HIV and SIV capsids

**DOI:** 10.1101/820886

**Authors:** Molly Ohainle, Kyusik Kim, Sevnur Keceli, Abby Felton, Ed Campbell, Jeremy Luban, Michael Emerman

## Abstract

The HIV-1 capsid protein makes up the core of the virion and plays a critical role in early steps of HIV replication. Due to its exposure in the cytoplasm after entry, HIV capsid is a target for host cell factors that act directly to block infection such as TRIM5 and MxB. Several host proteins also play a role in facilitating infection, including in the protection of HIV-1 capsid from recognition by host cell restriction factors. Through an unbiased screening approach, called HIV-CRISPR, we show that the Cyclophilin A-binding deficient P90A HIV-1 capsid mutant becomes highly-sensitized to TRIM5alpha restriction in IFN-treated cells. Further, the CPSF6-binding deficient, N74D HIV-1 capsid mutant is sensitive to restriction mediated by human TRIM34, a close paralog of the well-characterized HIV restriction factor TRIM5. This restriction occurs at the step of reverse transcription, is independent of interferon stimulation and limits HIV-1 infection in key target cells of HIV infection including CD4+ T cells and monocyte-derived dendritic cells. TRIM34 restriction requires TRIM5alpha as knockout or knockdown of TRIM5alpha results in a loss of antiviral activity. TRIM34 can also restrict some SIV capsids. Through immunofluorescence studies, we show that TRIM34 and TRIM5alpha colocalize to cytoplasmic bodies and are more frequently observed to be associated with infecting N74D capsids than with WT capsids. Our results identify TRIM34 as an HIV-1 CA-targeting restriction factor and highlight the potential role for heteromultimeric TRIM interactions in contributing restriction of HIV-1 infection in human cells.

## Introduction

The HIV-1 capsid protein (CA or p24gag) forms the core of the virion and is key to effective delivery of the HIV-1 genome inside a host cell and into the nucleus where integration into the host chromosome occurs (Campbell and Hope, 2015; Yamashita and Engelman, 2017). HIV-1 CA is involved in the early steps of HIV-1 replication including uncoating, nuclear entry, and integration site selection (Ambrose and Aiken, 2014; Campbell and Hope, 2015; Hilditch and Towers, 2014; Yamashita and Engelman, 2017). HIV-1 CA is also an important target for host restriction factors such as rhesus TRIM5alpha and MxB (Malim and Bieniasz, 2012; Yamashita and Engelman, 2017).

Many restriction factors are induced by type I Interferon (IFN). Recently, human TRIM5alpha, previously thought to lack activity against primate lentiviruses, has been shown to restrict wild type HIV-1 capsids in IFN-treated cells (Jimenez-Guardeno et al., 2019; OhAinle et al., 2018). TRIM5alpha restriction of retroviral capsids is driven by interactions between the C-terminal SPRY domain of TRIM5alpha and determinants present in assembled CA structures (Ganser-Pornillos and Pornillos, 2019). Although the affinity of the SPRY domain for CA is low, this low affinity is overcome by TRIM5alpha dimerization and its ability to form higher-order assemblies around the viral core, enhancing avidity of the TRIM5alpha-CA interaction (Ganser-Pornillos and Pornillos, 2019). TRIM5alpha is also able to oligomerize with other TRIM-family members (Li et al., 2011; Zhang et al., 2006). One key aspect of TRIM biology that remains relatively unexplored includes the potential for hetero-oligomerization of TRIM proteins that could have important functional consequences.

Through the study of HIV-1 CA mutants that lack binding to host cell factors or possess other key phenotypes, such as altered stability, much has been revealed about how CA determines the fate of HIV-1 cores inside cells. The host proteins CPSF6 and Cyclophilin A (CypA) have a complex but important role in HIV-1 CA interactions and infection (Yamashita and Engelman, 2017). HIV-1 CA binds CypA which provides protection against the action of TRIM5 (Kim et al., 2019; Luban et al., 1993). CPSF6 interacts with HIV-1 capsid on entry into target cells (Lee et al., 2010; Price et al., 2012) and facilitates interaction with nuclear import pathways that enhances targeting of HIV-1 integration into gene-rich regions (Rasheedi et al., 2016; Sowd et al., 2016). Single amino acid mutations in the HIV-1 capsid protein, for example N74D for CPSF6 and P90A for CypA, abrogate binding to these host factors (Lee et al., 2010; Schaller et al., 2011). Both capsid mutants have been demonstrated to infect cells less efficiently than wild type in some cell types, including primary cell such as CD4+ T cells and monocyte-derived macrophages (MDMs) (Ambrose et al., 2012; Bulli et al., 2016; Kim et al., 2019; Schaller et al., 2011). Further, both the P90A and N74D capsid mutants have been shown to be hypersensitive to the effects of IFN (Bulli et al., 2016), suggesting that one or more interferon-induced restriction factors block infection of these capsid mutant viruses. Restriction of these mutants has been shown to be independent of the IFN-induced capsid-targeting restriction factor MxB (Bulli et al., 2016) but identification of other capsid-targeting restriction factors underlying the increased IFN sensitivity of these CA mutants has been elusive.

Previously, we demonstrated that human genes that mediate the antiviral effects of IFN can be identified through an unbiased CRISPR screening approach called HIV-CRISPR (OhAinle et al., 2018). Here we use this approach to identify capsid-targeting restrictions that target the P90A and N74D HIV-1 capsid mutants. While the CypA-binding deficient P90A mutant becomes more sensitive to TRIM5alpha restriction, the CPSF6-binding deficient N74D mutant becomes sensitive to a novel restriction by the TRIM5alpha paralog, TRIM34. This restriction is independent of IFN induction as well as CPSF6 binding and results in a block during HIV reverse transcription. TRIM34 restriction occurs in primary cells in addition to the THP-1 monocytic cell line used in our screens. Further, we find that TRIM34 requires TRIM5alpha to inhibit the N74D while TRIM5alpha of the P90A virus occurs independent of TRIM34. Thus, we find that TRIM34 is a novel inhibitor of HIV-1 and SIV capsids that acts in conjunction with TRIM5 to limit infection of primary T cells.

## Materials and Methods

### Cells and Cell culture

All cells were incubated in humidified, 5% CO_2_ incubators at 37 °C. The THP-1 monocytic cell line (ATCC) was cultured in RPMI (Invitrogen) with 10% FBS, Pen/Strep, 10 mM HEPES, 0.11 g/L sodium pyruvate, 4.5 g/L D-Glucose and Glutamax. 293T (ATCC CRL-3216) and TZM-bl cells (ATCC 8129) were cultured in DMEM (Invitrogen) with 10% FBS and Pen/Strep. Puromycin selections in THP-1 cells were done at 0.5–1 ug/mL. The identity of THP-1 cells was confirmed by STR profiling (Fred Hutch Research Cell Bank). For the MoDC-related work, HEK293 cells (American Type Culture Collection) were cultured in DMEM supplemented with 10% heat-inactivated FBS, 20 mM GlutaMAX-I, 1 mM sodium pyruvate, 1× MEM non-essential amino acids and 25 mM HEPES, pH 7.2. HeLa and 293T cell lines utilized in the immunofluorescence assays were obtained from the American Type Culture Collection and were cultured in Dulbecco’s modified Eagle’s medium (DMEM) supplemented with 10% fetal bovine serum (FBS) (Atlanta Biologicals), 100 U/ml penicillin, 100 µg/ml streptomycin, and 10 µg/ml ciprofloxacin. To generate MoDCs, Peripheral Blood Mononuclear Cells (PBMCs) were isolated from leukopaks by gradient centrifugation on Lymphoprep (Axis-Shield Poc AS #AXS-1114546). CD14^+^ PBMCs were enriched using anti-CD14 antibody microbeads (Miltenyi Biotec #130-050-201), according to manufacturer’s protocol. The enriched CD14^+^ cells were cultured at a density of 2 × 10^6^ cells/mL, in RPMI-1640, supplemented with 5% heat-inactivated human AB^+^ serum (Omega Scientific), 20 mM GlutaMAX-I, 1 mM sodium pyruvate, 1× MEM non-essential amino acids and 25 mM HEPES pH 7.2. The addition of 1:100 cytokine-conditioned media containing hGM-CSF and hIL-4 to the culture promoted the differentiation of CD14+ cells into MoDCs. These cytokine-conditioned media were produced from HEK293 cells stably transduced with pAIP-hGMCSFco (Addgene no. 74168) or pAIP-hIL4-co (Addgene no. 74169), as previously described (McCauley et al., 2018; Pertel et al., 2011). For CD4+ T cell experiments, whole blood was obtained from BloodWorks Northwest, total PBMCs were isolated using the density gradient centrifugation method with Histopaque-1077 (Sigma-Aldrich #10771) and CD4+ T cells were isolated using EasySep Human CD4+ T cell isolation kit (StemCell Technologies #17952). Cells were resuspended to 2.5 × 10^6^ cells/mL in RPMI complete media supplemented with 10% FBS, Glutamax and Pen/Strep and with 100 U/mL recombinant human IL-2 (Roche; Sigma # 10799068001). For the analysis of reverse transcription products, a clonal TRIM34-KO THP-1 cell line was created through single-cell sorting of Cas9/RNP electroporated pools into 96-well plates to create individual clonal lines (BD FACS Aria II – Fred Hutch Flow Cytometry Core). A clonal KO line was identified through ICE Editing Analysis (Synthego). Universal Type I Interferon Alpha was obtained from PBL Assay Science (Catalog No. 11200– 2), diluted to 10^5^ Units/mL in sterile-filtered PBS/1% BSA according to the activity reported by manufacturer and frozen in aliquots at −80°C.

### Human blood

For monocyte-derived dendritic cell (MoDC) and activated CD4+ T cell preparations, leukopaks were acquired from anonymous, healthy blood donors (New York Biologics or BloodWorks Northwest). These experiments were declared to be non-human subjects research by the University of Massachusetts Medical School or Fred Hutchinson Cancer Research Center Institutional Review Boards, according to National Institutes of Health (NIH) guidelines (http://grants.nih.gov/grants/policy/hs/faqs_aps_definitions.htm).

### Plasmids

HIV infectious clones based on the LAI strain of HIV-1 (pBru3ori) were used in this study. The pBru3ori GFP3 backbone encodes the green fluorescent protein (GFP) gene in place of the *nef* gene (Yamashita and Emerman, 2004). The Bru3ori GFP3* WT, Bru3ori GFP3* N74D and Bru3ori GFP3* A77V proviruses were provided by Masahiro Yamashita and are described in (Saito et al., 2016). The P90A CA mutation was introduced into pBru3ori GFP3 using standard cloning procedures as described previously (Henning et al., 2014). The luciferase envelope-defective reporter proviral N74D plasmid was cloned from Bru3ori GFP3* N74D by BssHI and SalI digest and cloned into BruLuc2deltaEnv (Yamashita and Emerman, 2004). The SIVmacLUC E-R- and SIVagmLUC E-R-plasmids were a gift from Ned Landau (Mariani et al., 2003). The lentiCRISPRv2 plasmid was a gift from Feng Zhang (Addgene #52961). pMD2.G and psPAX2 were gifts from Didier Trono (Addgene #12259/12260). lentiCRISPRv2 constructs targeting genes of interest were cloned into BsmBI-digested lentiCRISPRv2 by annealing complementary oligos with overhangs that allow directional cloning into lentiCRISPRv2. TRIM34 oligos used were: TRIM34KO_1: TRIM34_1 Sense CACCGGTCAAGTTGAGCCCAGACAA and TRIM34_1 Antisense AAACTTGTCTGGGCTCAACTTGACC; TRIM34KO_2: TRIM34_2 Sense CACCGGAGTAACTGATACCACACAC and TRIM34_2 Antisense AAACGTGTGTGGTATCAGTTACTCC). TRIM5 oligos used were: TRIM5KO: TRIM5 Sense CACCGGTTGATCATTGTGCACGCCA and TRIM5 Antisense AAACTGGCGTGCACAATGATCAACC). The lentiviral pHIV-dTomato (Addgene #21374) expression vector was a gift from Bryan Welm (Addgene plasmid #21374; http://n2t.net/addgene:21374; RRID:Addgene_21374). The human TRIM34 cDNA was purchased from Genscript (NM_001003827.1). Human TRIM34 was cloned into pHIV/dTomato using NotI and XmaI sites with an HA tag encoded at the N-terminus. For HeLa immunofluorescence assays, YFP-TRIM5alpha and HA-TRIM34 were cloned in frame, into retroviral vectors EXN and YXN as described previously (Finzi et al., 1999).

### Virus and Lentivirus Production

Replication-competent HIV-1 viruses were produced as previously described (OhAinle et al., 2018). Briefly, 293T cells (ATCC) were plated at 2 × 10^5^ cells/mL in 2 mL in 6-well plates one day prior to transfection using TransIT-LT1 reagent (Mirus Bio LLC) with 3 μL of transfection reagent per μg of DNA. For HIV-1 production, 293Ts were transfected with 1 ug/well proviral DNA. One day post-transfection media was replaced. Two- or three-days post-transfection viral supernatants were clarified by centrifugation (1000 g) and filtered through a 20 μm filter. For Benzonase-treated viral preps, viral supernatants were incubated with 1 uL Benzonase (Sigma Aldrich #E1014) per 1mL of viral supernatant for 30 minutes at 37°C after dilution in 10X Benzonase Buffer (500mM Tris-HCl pH 8.0, 10mM MgCl_2_, 1 mg/mL Bovine Serum Albumin). For HIV-1 vectors used in Figure 4D-F, HEK293 cells were seeded at 75% confluency in 6-well plates. Transfections were performed with 6.25 μL TransIT LT1 transfection reagent (Mirus) in 250 μL Opti-MEM (Gibco) with 2.49 µg total plasmid DNA. 2.18 μg of env-defective HIV-1 provirus containing GFP reporter was cotransfected with 0.31 μg pMD2.G VSV G plasmid (Addgene #12259). Simian immunodeficiency virus (SIV)−VLPs containing Vpx were produced by the transfection of 2.18 μg pSIV−Δpsi/Δenv/ΔVif/ΔVpr (Addgene #132928) and 0.31 μg pMD2.G plasmid. 16 hours post transfection, the culture media was changed to the media for MoDC culture. Viral supernatant was harvested 2 days later, filtered through a 0.45 μm filter and stored at −80 °C. For lentiviral preps (lentiCRISPRv2 and pHIV), 293Ts were transfected with 667 ng lentiviral plasmid, 500 ng psPAX2 and 333 ng MD2G. For PIKA_HIV_ library preps, supernatants from 20 × 6 well plates were combined and concentrated by ultracentrifugation. 30 mL of supernatant per SW-28 tube were underlaid with sterile-filtered 20% sucrose (1 mM EDTA, 20 mM HEPES, 100 mM NaCl, 20% sucrose) and spun in an SW28 rotor at 23,000 rpm for 1 hr at 4°C in a Beckman Coulter Optima L-90K Ultracentrifuge. Supernatants were decanted, pellets resuspended in DMEM over several hours at 4°C and aliquots frozen at −80°C. All viral and lentiviral infections and transductions, except those in Figure 4D-F or Figure 5, were done in the presence of 20 μg/mL DEAE-Dextran (Sigma #D9885).

### HIV-CRISPR Screening & Screen Analysis

HIV-CRISPR Screening and Analysis was performed as described (OhAinle et al., 2018) with the ISG-specific PIKA_HIV_ library with the exception that the viral dose used in each screen allowed for infection of only ∼10-30% of cells for each capsid mutant virus. All screens were performed in a clonal THP-1 ZAP-KO cell line(OhAinle et al., 2018). Analysis of screen data was performed as previously described (OhAinle et al., 2018) with the exception that single mismatches were allowed when assigning reads to each sample during multiplexing.

### Transduction with lentiviral knockdown, knockout and overexpression vectors

For stable overexpression of TRIM34, THP-1 cells were transduced with pHIV/dTomato-TRIM34 or pHIV/dTomato empty vector lentiviral preps. 2 – 5 days post-transduction cells were sorted for high dTomato expression to select for high-expressing populations. Transduced cells were resorted as needed. For shRNA knockdown in MoDCs, 2 × 10^6^ CD14^+^ monocytes/mL were transduced with a 1:4 volume of SIV−VLPs and a 1:4 volume of knockdown lentivectors, as indicated. The SIV−VLPs were added to transfer Vpx to the cells in order to overcome restriction by SAMHD1 against lentiviral transduction (Hrecka et al., 2011; Laguette et al., 2011). Transduced cells were then selected with both 3 μg/mL puromycin (InvivoGen #ant-pr-1) and 10 μg/mL blasticidin (InvivoGen #ant-bl-1) for 3 days, starting at day 3 post-transduction. To generate stable HeLa cell lines used in immunofluorescence assays, a retrovirus was prepared by transfecting equal amounts of VSV-G, pCigB packaging plasmid, EXN HA-TRIM34 or YXN YFP-TRIM5 into HEK293T cells. Viral supernatant was harvested and filtered through 0.45 µm filters (Milipore) and applied to HeLa cells. 48 hrs after transduction, G418 was added to the cells, and following selection, cells were collected to check protein expression by Western blotting. To generate KO pools, THP-1 cells were transduced with lentiCRISPRv2 vectors and selection in Puromycin. KO cell pools were validated using genomic editing analysis as described below (Editing Analysis).

### Cas9/RNP Electroporation

Multiplexed Gene Knockout Kits targeting TRIM34 and TRIM5 were purchased from Synthego. The TRIM5 Kit includes the following sgRNA target sequences: AAUCUUGCUUAACGUACAAG, UGGCCACAGUCUAGACUCAA and GAGGCAGUGACCAGCAUGGG. Primers used to amplify the genomic locus and sequencing for TRIM5 were: GAAAAGCCCTTATTACCAGG (For) and GAGAATCCATGACTTGGAAG (Rev). The TRIM34 Kit includes the following sgRNA target sequences: AGGUCUUGUGGUUUGCAGUG, AGGGGUUAAUGUAAAGGAGG and GGGAACUGAUCCGGCACACA. TRIM34 amplification and sequencing primers were provided with the kit. CD4+ T cells were activated for 3 days with 10 ug/mL of plate-bound anti-CD3 (Tonbo Biosciences, clone UCHT1; #70-0038-U100) and 5 ug/mL of diffused anti-CD28 (Tonbo Biosciences, clone CD28.2; #70-0289-U100). For each electroporation 1 × 10^6^ CD4+ T cells were pelleted by centrifugation at 100 x *g* for 10 mins and washed once in PBS. Cells were resuspended in 25 uL of CRISPR/Cas9 crRNP complexes that were pre-assembled in P3 Primary Cell Nucleofector Solution (Lonza #V4SP-3096) before electroporation in a single well of a 96-well Nucleocuvette Plate using program EH-115. Each electroporation was diluted with 80 uL of RPMI complete + 125 U/mL IL-2 and allowed to recover for 1-2 hours at 37°C. Cells resuspended at 2.5 × 10^6^ cells/mL were transferred to a 96-well plate in RPMI complete + 100 U/mL IL-2 and re-activated with anti-CD2/CD3/CD28 beads at a 1:1 ratio (T Cell Activation/Expansion Kit, Milltenyi Biotec #130-091-441). Two days post-electroporation, each well was supplemented with 100 uL of RPMI complete + 100 U/mL IL-2. Beginning from 4 days post-electroporation, cells were maintained and propagated at 1 × 10^6^ cells/mL with fresh RPMI complete + 100 U/mL IL-2 being added every 2-3 days until infection and editing analysis.

### Editing Analysis

Cell populations were analyzed for allele editing frequency as previously described (OhAinle et al., 2018). Briefly, genomic DNA was isolated with a QIAamp DNA Mini Kit (Qiagen #51185), amplified by primers surrounding the editing site and sanger sequenced. Editing levels analyzed by ICE Analysis (Synthego) to obtain an ICE Editing Score.

### Exogenous reverse transcriptase assay

A 5 μL transfection supernatant containing virions was lysed in 5 μL 0.25% Triton X-100, 50 mM KCl, 100 mM Tris−HCl pH 7.4 and 0.4 U/µL RiboLock RNase inhibitor. This viral lysate was then diluted 1:100 in 5 mM (NH_4_)_2_SO_4_, 20 mM KCl and 20 mM Tris−HCl pH 8.3. 10 μL of this was then added to a single step, RT−PCR assay with 35 nM bacteriophage MS2 RNA (Integrated DNA Technologies) as a template, 500 nM of each primer (5’-TCCTGCTCAACTTCCTGTCGAG-3’ and 5’-CACAGGTCAAACCTCCTAGGAATG-3’) and 0.1 µL hot-start Taq DNA polymerase (Promega) in 20 mM Tris−HCl pH 8.3, 5 mM (NH_4_)_2_SO_4_, 20 mM KCl, 5 mM MgCl_2_, 0.1 mg/mL BSA, 1/20,000 SYBR Green I (Invitrogen) and 200 μM dNTPs in a total reaction volume of 20 μL. The RT−PCR reaction was carried out in a Bio-Rad CFX96 cycler with the following parameters: 42 °C for 20 min, 95 °C for 2 min and 40 cycles (95 °C for 5 s, 60 °C for 5 s, 72 °C for 15 s and acquisition at 80 °C for 5 s).

### Viral Infectivity Assays

Cells were pre-stimulated with IFNα 24 hr prior to infection where indicated. Virus and 20 μg/mL DEAE-Dextran in RPMI were added to cells, spinoculated for 20 min at 1100x*g*, and incubated overnight at 37°C. Cells were washed the next day and re-suspended in RPMI supplemented with IFNα. For infectivity assays using single-cycle viruses in MoDCs, 2.5 × 10^5^ cells were plated per well, in a 48-well plate, on the day of virus challenge. Media containing VSV G-pseudotyped HIV-1 vectors encoding GFP reporter was added to challenge cells in a total volume of 250 μL per condition. Cells were harvested for flow cytometric analysis by scraping at 48 hours post-challenge with the HIV-1 vectors and fixed in a 1:4 dilution of BD Cytofix Fixation Buffer with phosphate-buffered saline (PBS) without Ca2^+^ and Mg2^+^, supplemented with 2% FBS and 0.1% NaN3.

### Flow Cytometry

For intracellular Gag_p24_ (p24) staining, cells were harvested and fixed in 4% paraformaldehyde for 10 min and diluted to 1% in PBS. Cells were permeabilized in 0.5% Triton-X for 10 min and stained with 1:300 KC57-FITC (Beckman Coulter 6604665; RRID: <u>AB_1575987</u>). Data were collected on Accuri C6 (BD Biosciences – U Mass) or a BD FACSCANTO II (Fred Hutch Flow Cytometry Core) and analyzed with FlowJo software. For cell surface marker staining, cells were washed twice in PBS, stained in PBS/1% BSA, incubated at 4°C for 1 hr, washed twice in PBS, and analyzed on the Canto two flow cytometer (Fred Hutch Flow Cytometry Core).

### Luciferase assay

For analysis of Luciferase activity, infected cells were lysed in 100 μL BrightGlo Luciferase reagent (Promega #E2610) and read on a LUMIstar Omega Luminometer (settings: 1sec).

### qPCR Assay for HIV Late Reverse Transcription Products

For qPCR analysis of HIV late RT products, THP-1 cells were infected in 6-well plates with Benzonase-treated viral preparations. Total DNA was extracted from infected cells approx. 16 hours post-infection with a QIAprep Spin Miniprep kit (Qiagen #27106). HIV cDNA was amplified using TaqMan Gene expression Master Mix (AppliedBiosystems #4369016) with 900nM of each primer: J1 FWD (Late RT F) – ACAAGCTAGTACCAGTTGAGCCAGATAAG, J2 REV (Late RT R) – GCCGTGCGCGCTTCAGCAAGC and 250nM LRT-P (late RT Probe) – FAM-CAGTGGCGCCCGAACAGGGA-TAMRA. qPCR data was collected on an ABI QuantStudio5 Real Time (qPCR) System Instrument.

### Infection and Immunofluorescence Microscopy

HeLa cells were allowed to adhere to glass coverslips placed in wells of a 24-well plate. Synchronized infection was performed by spinoculation of reporter virus onto cells at 13 °C for 2 h at 1,200 × *g*, after which virus-containing medium was removed and replaced with warm media. Coverslips containing cells were incubated in 37 °C for 2 hours, and were subsequently fixed with 3.7% formaldehyde (Polysciences) in 0.1 M PIPES [piperazine-N,N’-bis(2-ethanesulfonic acid)], pH 6.8. Cells were permeabilized with 0.1% saponin, 10% normal donkey serum, 0.01% sodium azide in PBS. The following primary antibodies were used for immunofluorescence: mouse anti-HIV-1 p24 (Santa Cruz, Cat. # sc-69728), rabbit anti-HA (Sigma, Cat. # H6908). Primary antibodies were labeled with fluorophore-conjugated donkey anti-mouse or anti-rabbit antibody (Jackson ImmunoResearch Laboratories, Inc., West Grove, PA, USA). Images were collected with a DeltaVision microscope (Applied Precision, Issaquah, WA, USA) equipped with a digital camera (CoolSNAP HQ; Photometrics, Tucson, AZ, USA), using a 1.4-numerical aperture (NA) 100x objective lens, and were deconvolved with SoftWoRx software (Applied Precision, Issaquah, WA, USA).

### Image Analysis

For each condition in each experiment, 15 Z-stack images were acquired using identical acquisition parameters. Deconvolved images were analyzed using Imaris Software (Bitplane). For analysis of colocalization between p24 and HA-TRIM34 and/or YFP-TRIM5alpha, an algorithm was designed to create a three-dimensional surface around p24 signal. The algorithm was applied to all the images within a given experiment. Colocalization was defined as the presence of a signal intensity above a threshold value, and the same colocalization threshold was maintained for a given channel for all the images and conditions in a particular experiment. Both graphing and statistics calculations were performed in Prism (Graphpad Software, Inc).

## Results

### HIV-CRISPR screening identifies TRIM34 as an inhibitor of the N74D capsid mutant

The P90A and N74D capsid mutant viruses have been shown to be impaired in replication both in IFN-treated and untreated cells (Ambrose et al., 2012; Bulli et al., 2016; Rasaiyaah et al., 2013; Schaller et al., 2011). Therefore, we hypothesized that the P90A (CypA-deficient) and N74D (CPSF6-deficient) capsid mutants may be more sensitive to inhibition by capsid-targeting restriction factors in human cells. To identify the host cell restrictions targeting these capsid mutant viruses we used our unbiased screening approach, PIKA_HIV_ screening (OhAinle et al., 2018), to ask what Interferon-Stimulated Genes (ISGs) are responsible for inhibiting both mutants in THP-1 cells. HIV-CRISPR screening is a virus-packageable CRISPR screening approach in which infecting HIV virions package the HIV-CRISPR modified lentiviral vector *in trans* upon budding from the infected cell (OhAinle et al., 2018). As the level of virus replication is dependent on the phenotype of gene knockout introduced by Cas9 endonuclease and sgRNA encoded in the HIV-CRISPR vector, the virus itself serves to readout the barcodes of gene knockouts with effects on virus replication (Figure 1A). Quantification of individual 20bp sgRNA sequences enriched in the virions relative to the representation of sgRNA sequences in the cell populations allows for the identification of antiviral genes as gene knockouts that allow for more robust replication of each virus are enriched in the viral supernatant. In the HIV-CRISPR screens described here, a library of cells transduced with the HIV-CRISPR vector targeting Interferon-Stimulated Genes (ISGs), the PIKA_HIV_ library, was infected with wildtype or the N74D or P90A capsid mutant viruses after overnight treatment with Interferon. Two possibilities are that these mutants are either more sensitive to the same restrictions that target wild type capsids or that they are sensitive to novel capsid-targeting restriction factor(s).

**Figure 1.**
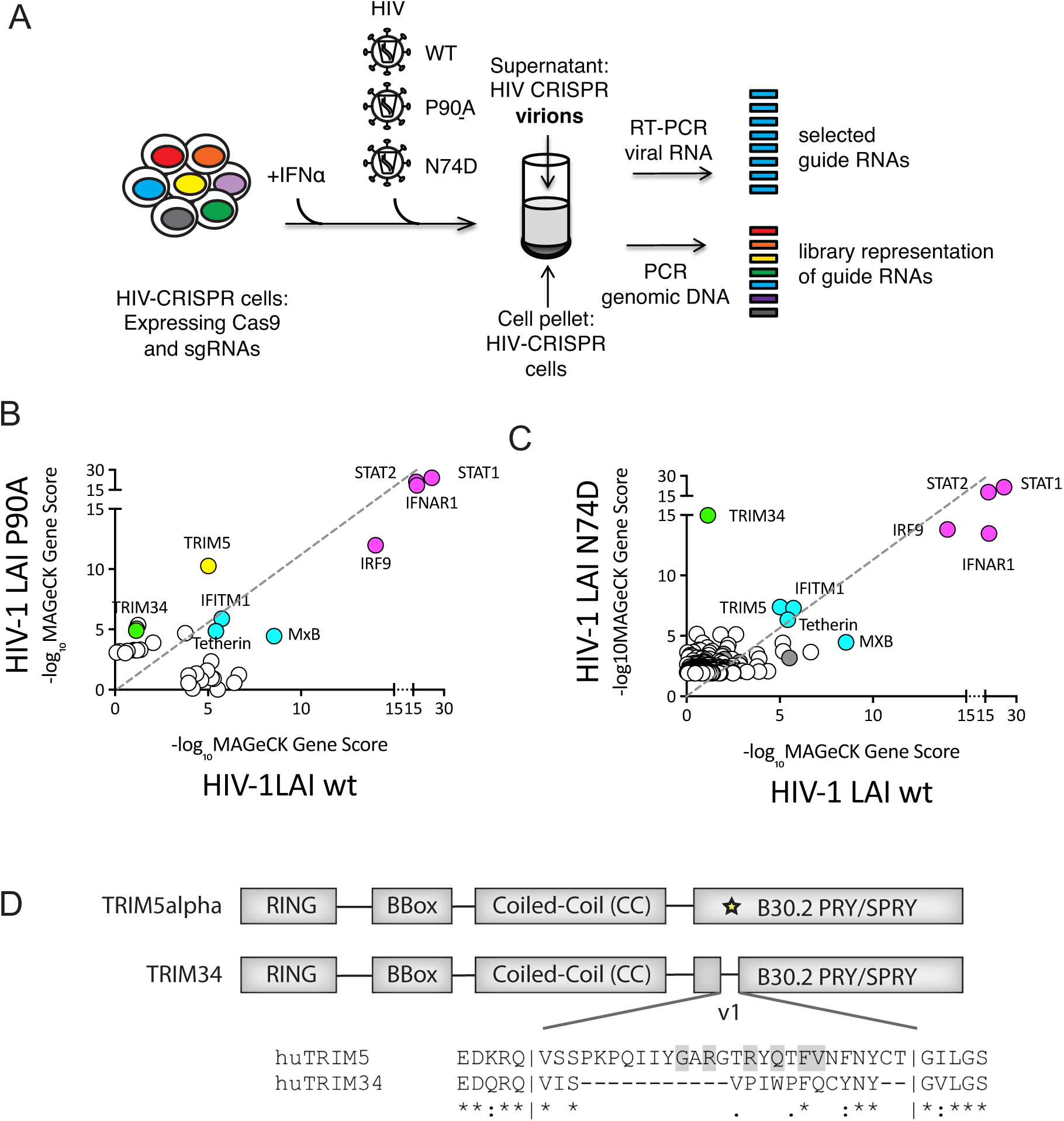
HIV-CRISPR screening identifies TRIM5alpha and TRIM34 as restriction factors for P90A and N74D HIV-1 CA mutants. <u>A</u>: HIV-CRISPR screens were performed after overnight IFN induction in THP-1 cells transduced with the PIKA_HIV_ (ISG-specific) library. HIV-CRISPR cells were infected with either wild type (n=2), the N74D CA mutant HIV-1_LAI_ (n=2), or the P90A CA mutant HIV-1_LAI_ (n=1). At 3 days post-infection cells and viral supernatants were collected, genomic DNA or viral RNA was extracted and 20bp sgRNA cassettes amplified by PCR or RT-PCR, respectively. <u>B and C</u>: MAGeCK Analysis of the enrichment of 20bp sgRNA sequences in viral RNA as compared to the genomic DNA was performed to calculate a MAGeCK Gene Score. Magenta: IFN pathway genes. Cyan: Gene hits shared across screens. Green: TRIM34. Yellow: TRIM5alpha. Gray: Non-Targeting Controls (NTCs) <u>B</u>: X-Axis: inverse log MAGeCK Gene Score for the WT HIV-1 screen. Y-Axis: inverse log MAGeCK Gene Score for the P90A HIV-1 CA mutant screen. The Top Gene hits are shown. <u>C</u>: X-Axis: inverse log MAGeCK Gene Score for the WT HIV-1 screen. Y-Axis: inverse log MAGeCK Gene Score for the N74D HIV-1 CA mutant screen. The Top Gene hits are shown. <u>D</u>: A schematic of the human TRIM5alpha and human TRIM34 domain structures, including the RING, B-Box, Coiled-Coil and B30.2 PRY/SPRY domains. An alignment of the v1 region of the SPRY domain (yellow star) is shown below. Gray shading indicates residues identified to be evolving under positive selection in Old World Monkey and Hominid TRIM5 by Sawyer et al (Sawyer et al., 2005).

Cell pellets and viral supernatants were collected and both genomic DNA from cells and viral RNA from virions were isolated and amplified for deep sequencing and MAGeCK analysis to identify sgRNA sequences significantly enriched in the viral supernatant of each virus. The WT and N74D screens were done in duplicate while the P90A screen represents a single replicate of the PIKA_HIV_ screen (Figure 1B: WT vs P90A; Figure 1C: WT vs N74D). The four genes in the library that are essential for IFN signaling, STAT1, STAT2, IFNAR1 and IRF9, were the highest-scoring hits in both screens as knockout of any gene in this pathway results in rescue of IFN inhibition of viral replication (Figure 1B and 1C: magenta). Next, a core set of ISGs, including MxB, IFITM1, Tetherin and TRIM5alpha, were some of the highest-scoring hits against wildtype, N74D and P90A viruses. Therefore, these restriction factors target the capsid mutants similar to what we previously found for wild type HIV-1 (OhAinle et al., 2018) and as we observed again here (Figure 1B and 1C: cyan). In both capsid mutant screens we measured a lower rank for MxB than for the wild type virus, consistent with the observed MxB resistance of both the P90A and N74D capsid mutants (Bulli et al., 2016) (Figure 1B and 1C – compare rank in WT screen as compared to either capsid mutant). In contrast, TRIM5alpha is the highest-scoring hit for the P90A virus (Figure 1B: yellow). This is consistent with recent results showing that loss of CypA binding by HIV-1 capsids results in sensitivity to TRIM5alpha restriction (Kim et al., 2019).

For the N74D capsid mutant screen we find a novel HIV-1 restriction factor, TRIM34, as the highest-scoring hit that is not found in the screen with wildtype HIV-1 (Figure 1C: Green). TRIM34 scores as highly as the IFN pathway genes (Figure 1C: Magenta), highlighting the key role TRIM34 plays in blocking replication of the N74D capsid mutant virus. TRIM34 was first described to be an Interferon-Stimulated Gene in HeLa cells (Orimo et al., 2000) and is in a cluster of paralogous human TRIM genes that includes TRIM5alpha, TRIM22 and TRIM6 on human chromosome 11 (Li et al., 2007). Like all members of the TRIM gene family (Ganser-Pornillos and Pornillos, 2019), TRIM34 encodes an N-terminal RBCC or tripartite motif, including RING, B-Box and coiled-coil domains (Figure 1D) (Orimo et al., 2000). TRIM34 shares 56% amino acid identity and overall domain structure with the closely-related, capsid-targeting HIV-1 restriction factor TRIM5alpha (Figure 1D). Therefore, TRIM34 could share some functional features of TRIM5alpha biology. However, unlike TRIM5alpha, TRIM34 has some surprising features. First, TRIM34 is not a significantly rapidly-evolving genes in primates (Li et al., 2007; Sawyer et al., 2007). Positive selection is frequently a feature of host antiviral genes in longstanding conflict with pathogens; therefore, the lack of positive selection in TRIM34 is unexpected. Further, human TRIM34 has a deletion in the v1 region of the B30.2 PRY/SPRY domain (Figure 1D), known to be important for determining the specificity of capsid recognition by TRIM5alpha (Sawyer et al., 2005). These findings suggest that while TRIM34 may have some structural and functional homology with TRIM5alpha, TRIM34 restriction of this HIV mutant is also likely to differ in some ways from TRIM5alpha.

### TRIM34 inhibits the N74D capsid mutant at a step before completion of reverse transcription

To validate that TRIM34 restricts the N74D capsid mutant virus but not wildtype HIV-1, we knocked out TRIM34 in THP-1 cells by transduction with a lentiviral vector encoding Cas9 and two different TRIM34-specific sgRNAs together with Non-Targeting Control (NTC) sgRNAs (Figure 2A and 2B). Efficient TRIM34 gene knockout was confirmed by sequencing analysis of the TRIM34 locus in both populations of cells (Figure 2A and 2B– 63% edited alleles for TRIM34KO_1 and 83% edited alleles for TRIM34KO_2). Control and TRIM34-KO cell populations were then infected with both WT and N74D capsid mutant viruses after overnight IFN treatment. We observe minimal effect of TRIM34 knockout on the replication of the wild type virus (Figure 2A). In contrast, we find 2.2-fold rescue of N74D viral replication in both TRIM34 KO pools (Figure 2A), consistent with our PIKA_HIV_ screen results that found TRIM34 as a significant block to infection in THP-1 cells.

**Figure 2.**
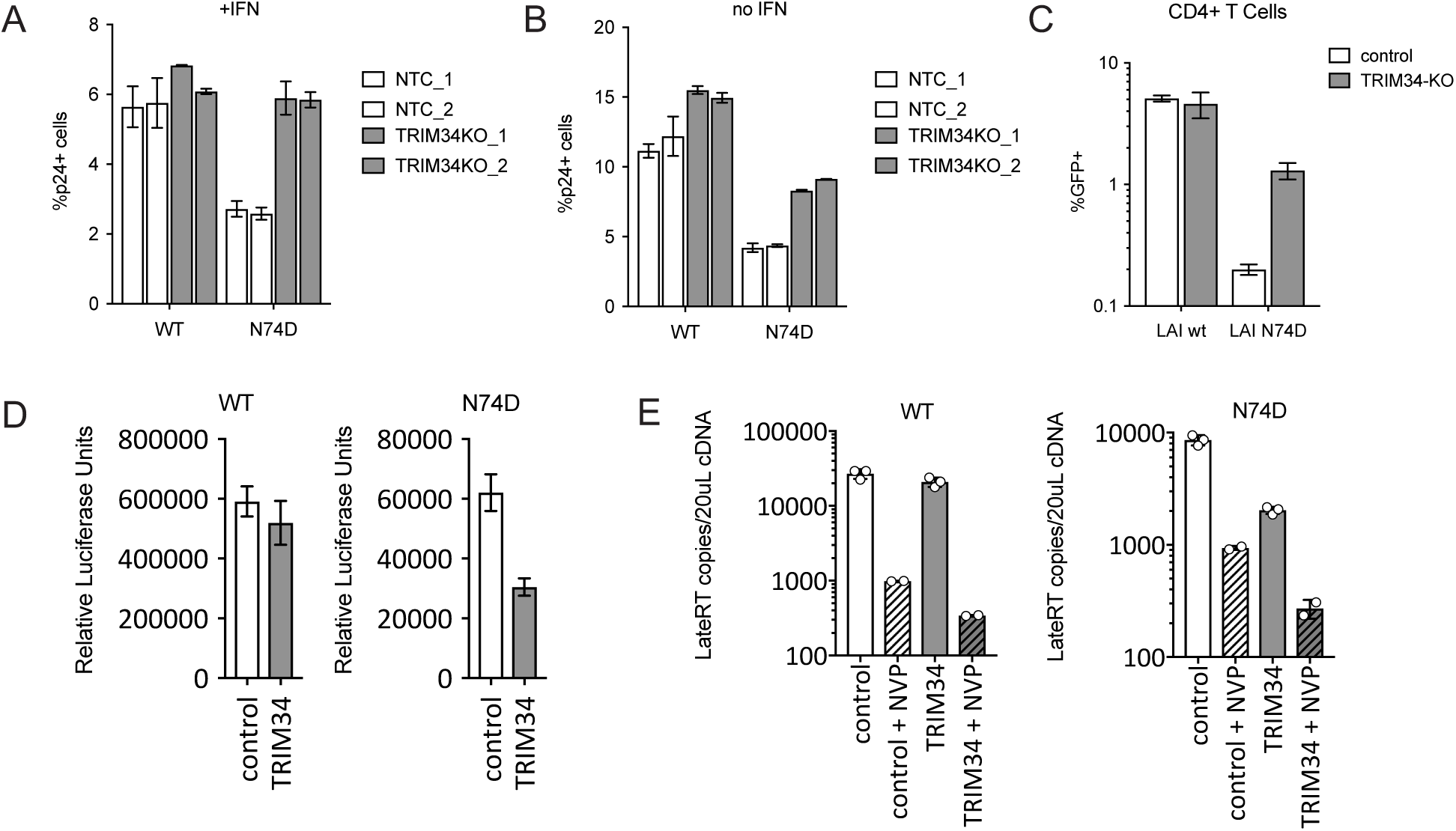
TRIM34 blocks HIV replication at the reverse transcription step in THP-1 and primary CD4+ T cells independent of IFN treatment. <u>A and B:</u> THP-1 cells were transduced with a lentiviral vector encoding Cas9 and an sgRNA targeting either TRIM34 (Gray Bars: n=2 pools, each an independent sgRNA) or a Non-Targeting Control (NTC) sgRNA (White Bars: n=2 pools, each an independent sgRNA). TRIM34 KO in each cell pool was determined by ICE Analysis (TRIM34KO_1 = 63% editing; TRIM34KO_2 = 83% editing. After overnight IFN stimulation cells were infected with either WT or N74D HIV and the amount of HIV replication assayed by staining for intracellular p24 2 days post-infection. Error bars indicate standard deviation from the mean from duplicate infections. <u>B:</u> Untreated cells (no IFN) were infected with either WT or N74D HIV and the amount of HIV replication assayed by staining for intracellular p24 2 days post-infection. Error bars indicate standard deviation from the mean from triplicate infections. <u>C:</u> Primary, activated CD4+ T cells were electroporated with Cas9-RNP complexes targeting TRIM34 (gray bars) or NTC (white bars). 2 days later edited CD4+ T cell pools were infected with GFP reporter HIV viruses (WT or N74D) and infection levels assayed 2 days later by flow cytometry. Error bars indicate standard deviation from the mean from triplicate infections. <u>D:</u> THP-1 cells were transduced with a lentiviral vector (pHIV) encoding TRIM34 (gray bars) or empty vector (white bars). Cell populations were sorted for dTomato expression and, following recovery, were infected with VSV-G pseudotyped WT or N74D Luciferase reporter viruses. Two days later levels of infection were assayed through a Luciferase Assay. Error bars indicate standard deviation from the mean from triplicate infections. <u>E:</u> TRIM34-overexpressing lines (gray bars) or control lines (white bars) were infected with WT-LUC or N74-LUC viruses with or without Nevirapine (NVP) to inhibit HIV reverse transcription. 16 hours later viral cDNA was collected and levels of HIV reverse transcription products were assayed by qPCR. Error bars indicate standard deviation from the mean from triplicate (no NVP) or duplicate (+NVP) infections as indicated.

To ask if TRIM34 restriction occurs only in IFN-treated cells as has recently been shown for TRIM5alpha (Jimenez-Guardeno et al., 2019), we also infected control and TRIM34-KO cells with wildtype and N74D virus without any IFN treatment (Figure 2B). Unlike TRIM5alpha inhibition of wild type HIV-1 capsids, TRIM34 is a constitutive inhibitor of the N74D CA mutant as KO rescues infection of the N74D virus even in the absence of IFN treatment 2-Fold (Figure 2B). Further confirming these findings, we repeated the PIKA_HIV_ screen in THP-1 cells without any IFN treatment and find TRIM34 as a significant hit (data not shown).

We also determined whether or not TRIM34 plays a role in restriction of HIV-1 capsids in primary activated, CD4+ T cells. TRIM34 KO CD4+ T cell pools were generated through electroporation of CD3/CD28-activated CD4+ T cells with CRISPR/Cas9 complexes targeting TRIM34 or NTC. 62% of TRIM34 alleles were found to be edited in this experiment (Figure 2C). We infected the control and TRIM34-KO pools with both wild type and the N74D capsid mutant viruses and measured infection levels after 2 days by flow cytometry. Consistent with our results in the THP-1 cell line model, knockout of TRIM34 rescues N74D CA infection 6.5-fold in CD4+ T cells (Figure 2C). Therefore, TRIM34 is endogenously-expressed and functions as an HIV-1 restriction factor in these key HIV-1 target cells.

TRIM5alpha binds to and blocks HIV-1 capsids in the cytoplasm, resulting in significantly decreased reverse transcription products during infection (Ganser-Pornillos and Pornillos, 2019). Moreover, there is a block prior to reverse transcription of the N74D CPSF6-binding capsid mutant in macrophages (Ambrose et al., 2012). Thus, we hypothesized that TRIM34 would mediate a similar block before reverse transcription would occur. In order to test this hypothesis, we stably-overexpressed human TRIM34 in clonal TRIM34-KO THP-1 cells and assayed replication of both the wild type and the N74D CA mutant viruses. TRIM34 overexpression does not result in any inhibition of wildtype HIV-1 (Figure 2D, left panel). In contrast, we observe significant inhibition of the N74D mutant by TRIM34 (Figure 2D, right panel). Therefore, overexpression of TRIM34 in THP-1 cells does allow for restriction of the N74D CA mutant virus. To ask if TRIM34 blocks infection before completion of reverse transcription, similar to rhesus TRIM5alpha, we infected control and TRIM34-overexpressing THP-1 cells and assayed viral DNA accumulation through a qPCR assay that detects HIV-1 reverse transcription products (De Iaco and Luban, 2011). The inhibition of replication of the N74D CA virus (Figure 2D) in TRIM34-overexpressing cells is correlated with a similar decrease in the accumulation of HIV viral DNA (Figure 2E). Therefore, TRIM34, like its paralog TRIM5alpha, inhibits HIV-1 replication early in the viral life cycle before the completion of reverse transcription.

### TRIM34 restricts HIV-1 and SIV capsids independent of CPSF6 binding

The N74D CA mutant virus was first characterized due to its loss of CPSF6 binding (Lee et al., 2010). Therefore, we reasoned that loss of CPSF6 binding could expose HIV-1 CA to restriction by TRIM34. To ask if the loss of CPSF6 binding is sufficient to sensitize HIV-1 capsids to TRIM34 restriction, we tested another CPSF6-binding capsid mutant A77V that, like N74D, also results in loss of binding to CPSF6 (Saito et al., 2016). In contrast to infection with N74D CA we find that infection of TRIM34-overexpressing cells is equivalent to wild type cells for the A77V mutant (Figure 3A). We further tested the relative sensitivity of N74D and A77V in primary cells by knocking out TRIM34 in primary, activated CD4+ T cells (Figure 3B). We measured infection of these cells by the N74D and A77V CA mutants as compared to a wild type control (Figure 3B). As shown in Figure 2, infection with the N74D CA mutant can be rescued by TRIM34 knockout in CD4+ T (Figure 3B). Consistent with our overexpression assay in THP-1 cells we find that the A77V mutant is not rescued by TRIM34 knockout in CD4+ T cells (Figure 3B). Therefore, in both THP-1 cells and primary CD4+ T cells the A77V mutant that lacks binding to CPSF6 does not become sensitive to TRIM34 restriction. Therefore, restriction by TRIM34 is independent of CPSF6 binding status of the HIV-1 capsid. Instead restriction of the N74D CA by TRIM34 is determined by a feature of this capsid other than CPSF6 binding.

**Figure 3.**
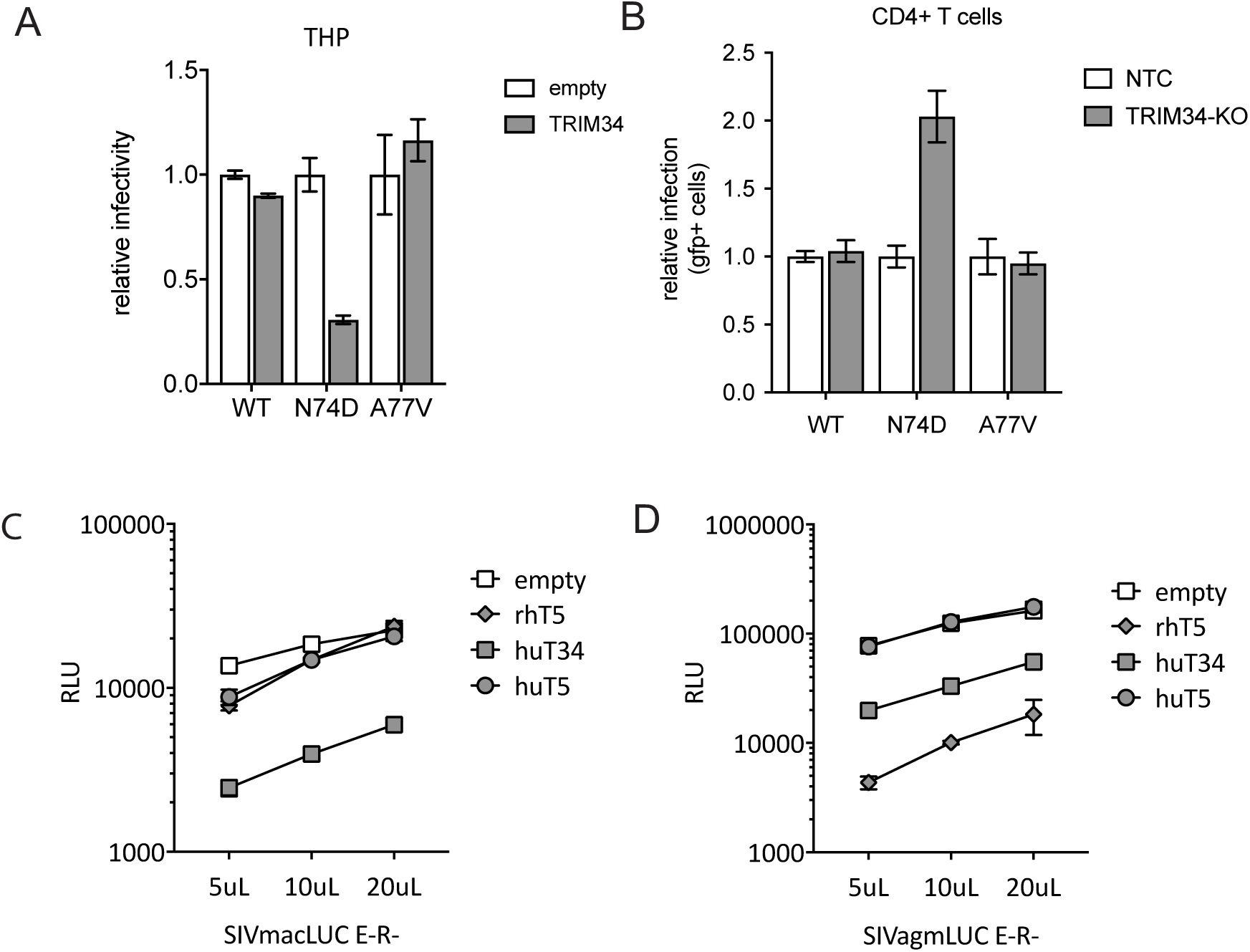
TRIM34 inhibits a range of primate lentiviruses independent of CPSF6. <u>A:</u> THP-1 cells stably-overexpressing TRIM34 (gray bars) or control cells (white bars) were infected with WT, N74D or A77V HIV-1 and levels of infection assayed 2 days post-infection by flow cytometry. The relative infection is normalized to the average infection in the control cells for each virus. Error bars indicate standard deviation from the mean from triplicate infections. <u>B:</u> Primary, activated CD4+ T cells were electroporated with Cas9-RNP complexes targeting TRIM34 (gray bars) or control crRNAs (white bars: control). 2 days later edited CD4+ T cell pools were infected with GFP reporter HIV viruses (WT, N74D or A77V) and infection levels assayed 2 days later by flow cytometry. Error bars indicate standard deviation from the mean from triplicate infections. <u>C and D:</u> THP-1 cells were transduced with lentiviral vectors encoding rhesus TRIM5alpha (gray diamonds: rhT5), human TRIM5alpha (gray circles: huT5), human TRIM34 (gray squares: huT34) or a control vector (white squares: empty). Each cell pool was infected with VSV-G pseudotyped SIVmacLUC (C) or SIVagmLUC (D) at 3 viral doses as indicated and levels of infectivity assayed 2 days later by luciferase assay. Error bars indicate standard deviation from the mean from triplicate infections.

Restriction factors from one species often potently restrict primate lentiviruses adapted to replicate in other primates to due to ongoing genetic conflict between hosts and pathogens (Duggal and Emerman, 2012). To ask if human TRIM34 may restrict lentiviruses more broadly, including Simian Immunodeficiency Viruses (SIVs) that are adapted to infect Old World Monkeys, we assayed human TRIM34 for restriction of SIVagm and SIVmac in THP-1 cells. In contrast to human or rhesus TRIM5alpha, human TRIM34 overexpression inhibits replication of SIVmac significantly (Figure 3C). Similarly, replication of SIVagm is significantly blocked by human TRIM34 overexpression, although this restriction is not as potent as rhesus TRIM5alpha restriction of this virus (Figure 3D). Therefore, human TRIM34 retrovirus restriction activity is not limited to HIV-1 CA mutants, as TRIM34 restricts at least two SIV strains in addition to the HIV-1 N74D CA mutant. Moreover, because SIVmac is restricted by TRIM34, yet is known to bind CPSF6 (Lee et al., 2010), this further supports the model that TRIM34 restriction sensitivity is not necessarily linked to CPSF6 binding of HIV or SIV capsids.

### TRIM34 restriction requires TRIM5alpha

TRIM34 is a close paralog of TRIM5alpha which is known to dimerize and form higher-order oligomers (Ganser-Pornillos and Pornillos, 2019). Furthermore, TRIM34 has been shown to be able to interact with TRIM5alpha in cells both in a yeast two-hybrid assay (Zhang et al., 2006) as well as in immunoprecipitation studies (Li et al., 2007; Li et al., 2011). As TRIM34 lacks a signal of positive selection in the B30.2 PRY/SPRY domain (Li et al., 2007; Sawyer et al., 2007), we reasoned that the association with TRIM5alpha may give TRIM34 specificity for HIV capsids. In this case, TRIM34 would require TRIM5alpha to restrict HIV-1 infection. This model is particularly intriguing as the PRY/SPRY domain of TRIM34 has a deletion in the v1 loop (see Figure 1D), a region shown to be critical for mediating specificity of capsid recognition by TRIM5alpha (Sawyer et al., 2005).

To ask if TRIM5alpha is important for restriction of the N74D capsid mutant virus, we compared the ability of TRIM34 to restrict the N74D virus in cells both with and without TRIM5alpha. We introduced a TRIM5alpha or Control CRISPR/Cas lentiviral vector into THP-1 cells overexpressing TRIM34 (Figure 4A). Infection of these TRIM34-overexpressing, TRIM5alpha-KO THP-1 pools demonstrates that the restriction of viral replication measured for the N74D capsid mutant is lost when TRIM5 is missing (Figure 4A). In contrast, we see no effect of TRIM34 overexpression or TRIM5-KO in THP-1 cells infected with the WT capsid (Figure 4B), consistent with there being no role for restriction of WT capsid by either TRIM34 or TRIM5alpha in cells lacking IFN stimulation. Therefore, TRIM34 is dependent on TRIM5alpha for full restriction activity.

**Figure 4.**
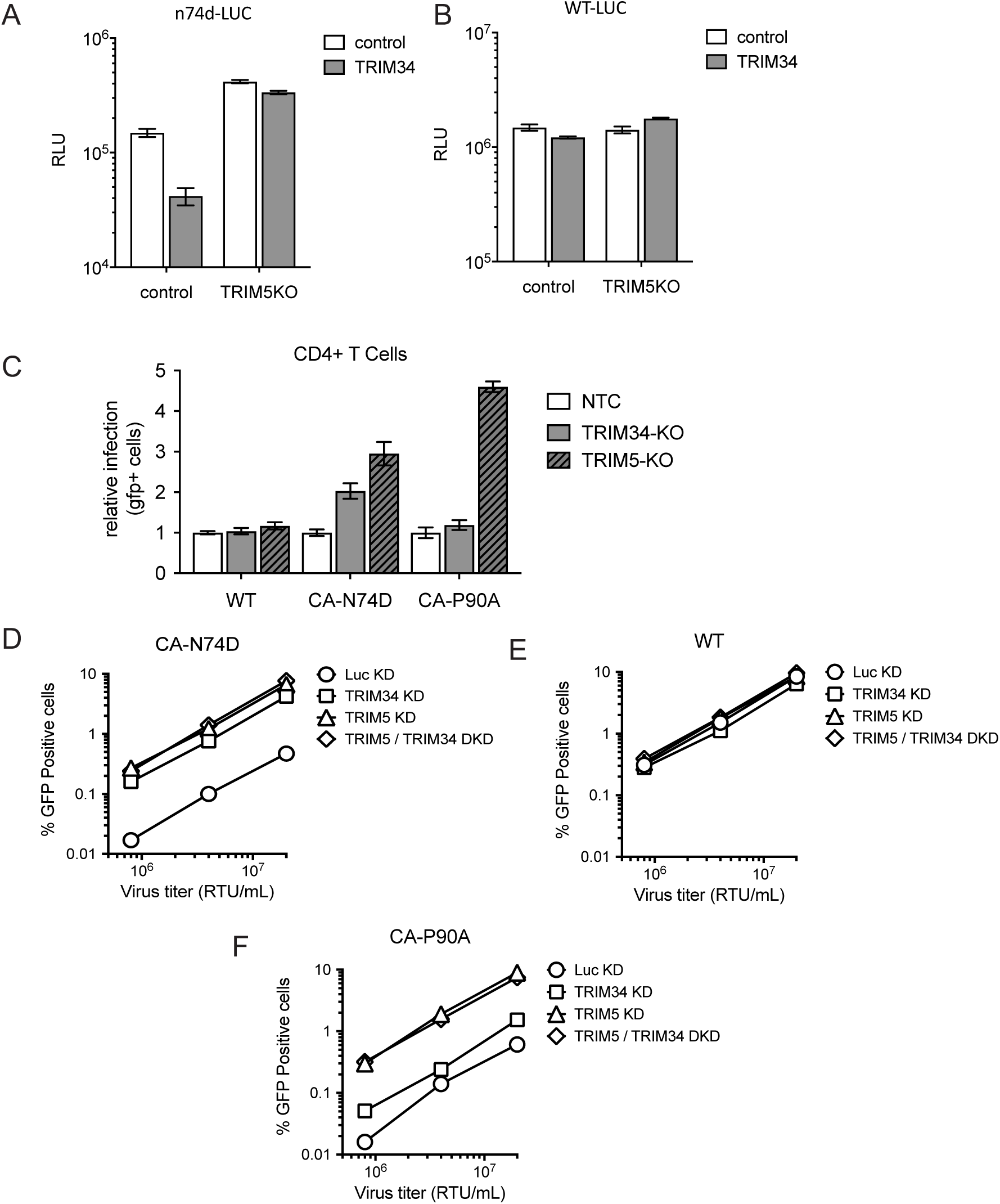
TRIM34 requires TRIM5alpha to restrict the N74D virus. <u>A and B:</u> THP-1 cells stably-overexpressing TRIM34 (gray bars) or control cells (white bars) were transduced with sgRNA-encoding lentiviral vectors targeting TRIM5alpha (TRIM5KO) or a control. TRIM5 alleles were edited at 70% for both control and TRIM34-overexpressing cells as determined by ICE analysis. Cell pools were infected with the N74D-LUC virus (A) or WT-LUC virus (B) and levels of infection assayed 2 days post-infection by luciferase assay. Error bars indicate standard deviation from the mean from triplicate infections. <u>C:</u> Primary, activated CD4+ T cells were electroporated with Cas9-RNP complexes targeting TRIM34 (gray bars), TRIM5alpha (hatched, dark gray bars) or NTC crRNAs (white bars). 2 days later edited CD4+ T cell pools were infected with GFP reporter HIV-1 viruses (WT, N74D or P90A) and infection levels assayed 2 days later by flow cytometry. The relative infection is normalized to the average infection in the control cells for each virus. Error bars indicate standard deviation from the mean from triplicate infections. <u>D, E and F:</u> Monocyte-derived dendritic cells were simultaneously transduced with a lentiviral vector encoding shRNAs targeting TRIM34 or luciferase control (Luc) and the other shRNA vector specific for TRIM5 or Luc, for the knockdown as indicated. The pooled cells were challenged with VSV-G pseudotyped HIV-1 vectors expressing GFP reporter and containing CA-N74D (D), WT CA (E), or CA-P90A (F), across a range of viral inputs. The percentage of GFP-positive cells was determined 2 days later by flow cytometry.

To ask if this requirement of TRIM5alpha for restriction of the N74D virus by TRIM34 is also important in primary cells, we knocked out either TRIM34 or TRIM5alpha in primary CD4+ cells and infected each cell pool with WT, N74D or P90A CA mutant viruses in comparison to a control cell pool (Figure 4C). Indeed, consistent with a requirement of TRIM5 for the TRIM34-medicated restriction, we find that knockout of either TRIM34 or TRIM5 is sufficient to rescue infection with the N74D capsid mutant (Figure 4C) while neither knockdown has a significant effect on WT HIV-1 infection (Figure 4C). We also examined this question in primary monocyte-derived dendritic cells (MoDCs) by transducing them with shRNA lentiviral vectors, resulting in stable knockdown of TRIM34 (TRIM34 KD) or TRIM5alpha (TRIM5 KD) (Figure 4D). In addition, to more directly ask if TRIM34 and TRIM5alpha work together or synergistically, we challenged cells in which both TRIM34 and TRIM5alpha were depleted simultaneously (TRIM5/TRIM34 DKD). Similar to our results in THP-1 and primary CD4+ T cells, infection with the N74D capsid mutant can be rescued by knockdown of either TRIM34 or TRIM5alpha in MoDCs (Figure 4D).

In contrast, knockdown of either TRIM34 or TRIM5alpha does not have any effect on wild type HIV-1 capsids (Figure 4E). Further, TRIM34 and TRIM5alpha act in the same pathway, rather than synergistically, to inhibit the N74D capsid mutant since a double-knockdown in MoDCs does not show any additional rescue (Figure 4D).

Finally, we asked if TRIM5alpha restriction of the CypA-binding deficient P90A mutant depends on TRIM34. Consistent with the results of our initial screen with the P90A mutant (Figure 1B), we find that HIV-1 P90A is sensitive to TRIM5alpha restriction (Figure 4C: CD4+ T Cells; Figure 4F: MoDCs). However, this restriction activity is independent of TRIM34 as TRIM34 knockout or knockdown has little to no effect on the P90A virus (Figure 4C: CD4+ T Cells; Figure 4F: MoDCs). These data suggest that TRIM34 restriction depends on TRIM5alpha, but that TRIM5alpha restriction does not depend on TRIM34. Therefore, there is asymmetry in the TRIM5alpha/TRIM34 relationship as their interdependence is not equivalent across restriction activities.

### TRIM34 and TRIM5alpha complexes colocalize with the N74D capsid mutant

Since TRIM34 restriction depends on TRIM5 (Figure 4), we tested the hypothesis that TRIM34 and TRIM5 directly colocalizes with each other and with incoming HIV-1 capsids during infection. HeLa cell lines stably-expressing YFP-TRIM5 and HA-TRIM34 were infected with WT HIV-1 or the N74D CA mutant and 2 hours later colocalization of TRIM34 or both TRIM34 and TRIM5alpha with each capsid was compared. Given that TRIM34 restricts N74D capsids but not the WT virus, we hypothesized that this differential restriction could be due to specific localization of TRIM34 to N74D HIV-1 capsids that does not occur, or does not occur to the same extent, as with wild type HIV-1 capsids. TRIM34 localizes to cytoplasmic puncta, commonly referred to as cytoplasmic bodies, similar to TRIM5alpha (Figure 5A – “Mock”; red – TRIM34, green – TRIM5alpha). Indeed, we observed colocalization of TRIM34 with HIV-1 capsids in the cell cytoplasm (Figure 5A: compare “WT” to “N74D” – white arrowheads). Quantification of the number of p24+ puncta that are also positive for TRIM34 shows that colocalization of TRIM34 with p24 occured more frequently for N74D capsids than for WT capsids (Figure 5B). As TRIM34 restriction of the N74D capsid mutant requires TRIM5alpha, we also quantified the degree to which TRIM34 colocalizes with TRIM5alpha in cells. We assayed the subcellular localization of both TRIM34 and TRIM5alpha in HeLa cells stably-overexpressing both YFP-TRIM5alpha and HA-TRIM34. We observe significant colocalization of TRIM34 and TRIM5alpha in the same cytoplasmic bodies (Figure 5A “Mock”: yellow – white triangles). Further, complexes containing both TRIM34 and TRIM5alpha preferentially colocalize with the N74D capsid mutant cores as compared to WT cores (Figure 5C). Therefore, TRIM34 and TRIM5 are present together with incoming N74D HIV-1 capsids in the cytoplasm of infected cells.

**Figure 5.**
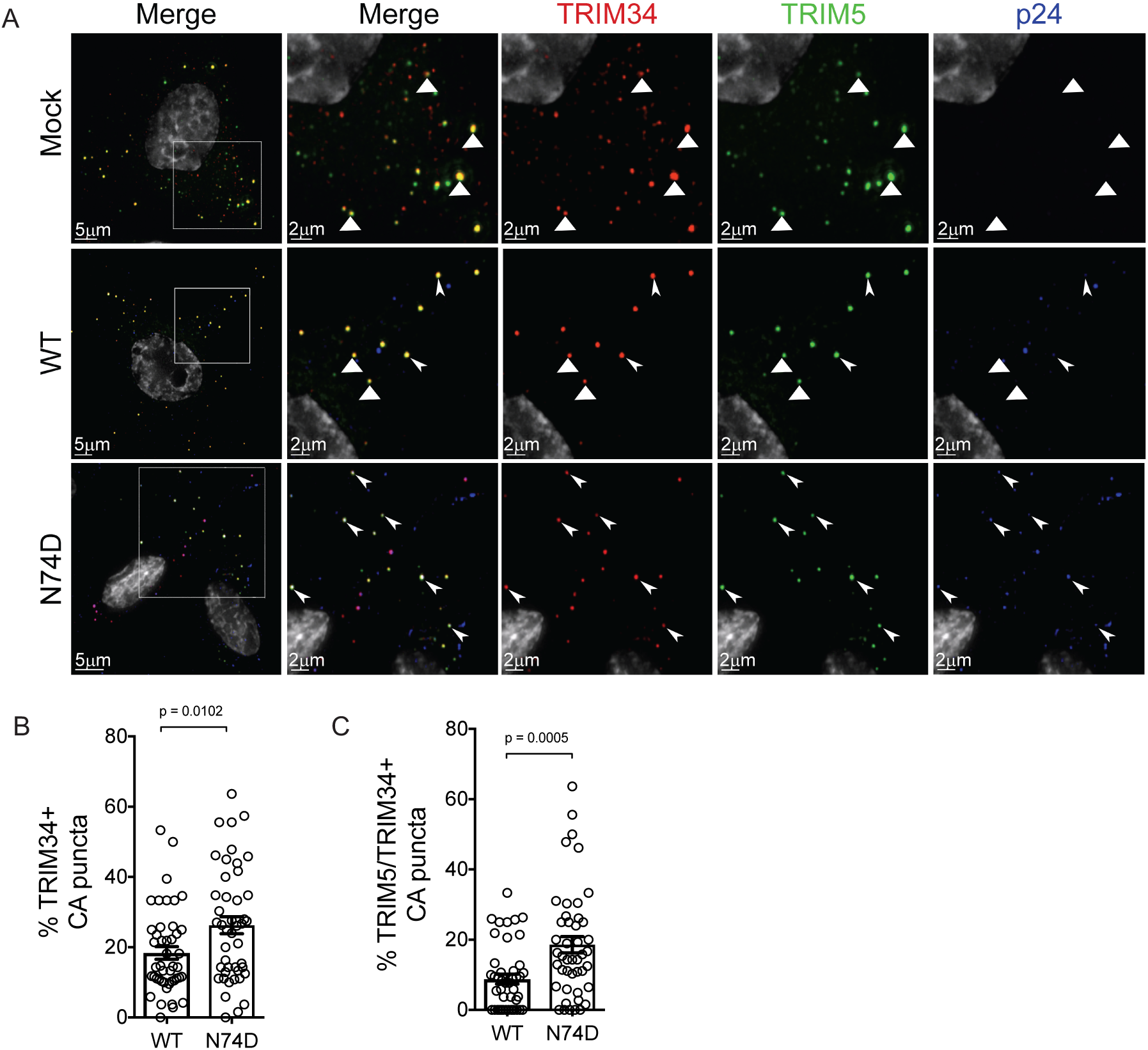
TRIM34 colocalizes more frequently with the restricted HIV-1 N74D capsid. HeLa cells were transduced to express YFP-TRIM5alpha (green) and HA-TRIM34 stably. They were plated on coverslips and synchronously infected with VSV-G pseudotyped HIV-1 with WT or N74D capsids as indicated. At 2 hpi, cells were fixed and stained for viral capsid protein p24 (blue) and HA-TRIM34 tag (red). 15 images were collected per condition in three independent biological replicates. <u>A:</u> Representative images for mock-infected cells (top row), WT-infected cells (middle row), and N74D-infected cells (bottom row). Areas of colocalization between TRIM34 and TRIM5 are indicated by triangles, and triple colocalization between TRIM24, TRIM5 and p24 are indicated by arrows in the zoomed in images for each channel. <u>B:</u> Quantification of percent p24 colocalizing with TRIM34 for the WT and N74D virus. *P* value was determined by an unpaired t test. Error bars represent SEM of all images collected across three biological replicates for each condition. <u>C:</u> Quantification of percent p24 colocalizing with both TRIM34 and TRIM5alpha for the WT and N74D virus. p value was determined by an unpaired t test. Error bars represent of all images collected across three biological replicates for each condition.

## Discussion

We identified TRIM34, a TRIM5 paralog, as an HIV-1 restriction factor capable of inhibiting infection by the N74D capsid mutant virus as well as several lentiviruses from monkeys. This block occurs before the completion of reverse transcription and is a constitutive block to infection as the restriction is observed in both IFN-stimulated and unstimulated cells. The antiviral activity of TRIM34 is independent of the ability of viral capsids to bind CPSF6 as shown with the CPSF6-binding deficient capsid mutant, A77V, that is not restricted by TRIM34. Moreover, TRIM34 restriction also occurs in primary, activated CD4+ T cells and monocyte-derived dendritic cells (MoDCs). The antiviral restriction activity of TRIM34 requires TRIM5alpha in all cells tested, however restriction of the HIV-1 P90A capsid mutant by TRIM5alpha does not require TRIM34. Finally, TRIM34 and TRIM5 colocalize in cells and preferentially localize to restricted N74D HIV-1 capsids as compared to WT capsids.

### TRIM34 is a capsid-targeting HIV-1 restriction factor

Our HIV-CRISPR screen demonstrates that the increased IFN-sensitivity of the N74D CA mutant (Bulli et al., 2016) is not due to a shift in sensitivity to the known capsid-targeting restriction factors TRIM5alpha or MxB (Figure 1B) but rather is due to restriction by a novel capsid-targeting restriction factor, TRIM34. TRIM34 is a close paralog of the well-studied capsid-targeting restriction factor TRIM5alpha. Previously, modest restriction of SIVmac by human TRIM34 was described but no effect of TRIM34 was observed on either wild type HIV-1 or a G98V capsid mutant virus (Zhang et al., 2006). Our data suggest that TRIM34, together with TRIM5alpha, mediates all or most of the block to replication of the N74D capsid mutant observed in primary cells, including CD4+ T cells and MDMs, first described by Ambrose et al. (Ambrose et al., 2012).

The site that sensitizes HIV-1 capsid to TRIM34 restriction, N74, is highly-conserved across HIV-1 and SIV strains (Lee et al., 2010). Mutations at this site may sensitize capsids to TRIM34/TRIM5 restriction and therefore have been selected against. However, this effect is independent of CPSF6 binding to capsids as loss of binding to CPSF6 is not sufficient for restriction by TRIM34 (see the A77V mutant in Figure 3A and 3B). Our results are consistent with the finding that wild type capsids do not become IFN-hypersensitive in CPSF6 knockout cells (Bulli et al., 2016). Therefore, our data support a model in which CPSF6 binding plays a role in integration targeting (Achuthan et al., 2018; Bejarano et al., 2019; Rasheedi et al., 2016; Schaller et al., 2011; Sowd et al., 2016) but it does not shield HIV-1 capsids from capsid-targeting restrictions. In contrast, loss of CypA binding sensitizes HIV-1 capsids to TRIM5alpha restriction (Kyusik’s paper). However, loss of CypA binding is not sufficient for TRIM5alpha restriction of all primate lentiviruses as SIVmac is not restricted by TRIM5alpha even though SIVmac capsids do not bind CypA to any appreciable affinity (Schaller et al., 2011). These data together highlight the complexity of capsid adaptation to host restrictions where multiple, independent pathways of adaptation to multiple restrictions may occur across divergent strains.

The capsid-targeting restriction factors MxB and TRIM5alpha together with other restriction factors likely constrain HIV-1 evolution *in vivo* and provide a significant host barrier that HIV-1 must adapt to in order to successfully establish infection during transmission to a new host (Asmuth et al., 2010). Capsid is a dominant determinant of species tropism for cross-species replication of primate lentiviruses such as SIVmac (Hatziioannou et al., 2006). Our results suggest that TRIM34, like TRIM5alpha, is a lentiviral restriction factor that HIV-1 has adapted to in order to replicate efficiently in human cells since SIVs are also sensitive to human TRIM34.

TRIM5alpha knockout or knockdown rescues the HIV-1 P90A capsid mutant in human primary cells suggesting that at least one function of CypA binding is evasion of TRIM5alpha-mediated restriction (Kyusik’s paper here). Our screening data show with the P90A mutant did not uncover an additional factor other than TRIM5, suggesting that the increased IFN sensitivity of this mutant is (Jimenez-Guardeno et al., 2019) is entirely due to TRIM5 induction by IFN.

### TRIM34 restriction requires TRIM5alpha

Of particular interest, we find that TRIM34 restriction depends on TRIM5alpha. More broadly, TRIM proteins are a large gene family (Nisole et al., 2005) and potentially-important functional interactions between family members likely have been overlooked. Our data show that TRIM34 can restrict lentiviral capsids but that this activity requires expression of TRIM5alpha, a close paralog of TRIM34. The ability of TRIM family members to hetero-oligomerize has been known for some time, but our screen highlights the power of an unbiased approach such as HIV-CRISPR screening to uncover previously-unappreciated functional interactions between TRIM family members.

This dependence of TRIM34 on TRIM5alpha raises several key questions including: 1) how does TRIM34 allow for specific restriction of the N74D capsid mutant viruses? and 2) why does this restriction depend on TRIM5alpha? We propose two potential models that are consistent with our data that are not mutually exclusive. First, it is possible that TRIM34 changes the specificity of TRIM5alpha such that it can now recognize the N74D capsid mutant virus better and/or more efficiently. TRIM34 can bind to HIV-1 capsids through its SPRY domain (Li et al., 2007; Yang et al., 2014). However, when compared to TRIM5alpha, human TRIM34 has a deletion in the v1 loop of the SPRY domain that is important for determining capsid binding specificity. Further, TRIM34 is not itself evolving under positive selection, an evolutionary signature that would be consistent with a direct protein interface with a viral pathogen. Instead, it may be that the TRIM34 SPRY does not itself make significant contact with the N74D capsid but that TRIM34 complexing with TRIM5alpha may affect recognition of this capsid specifically by human TRIM5alpha.

A second possibility is that TRIM34 stabilizes TRIM5alpha, thereby allowing it to restrict the N74D capsid mutant virus. This model does not require any change in specificity of binding to capsid, *per se*, but rather results from changes in kinetics and/or entry of HIV-1 capsid mutants. The N74D has the same intrinsic stability as the wild type capsid as measured by an *in vitro* uncoating assay (Shah et al., 2013). Further, the capsid stability and kinetics of uncoating of the N74D virus have been reported to be similar to WT (Shah et al., 2013). In contrast, slower uncoating kinetics of the N74D capsid mutant has been observed (Hulme et al., 2015). In support of this model, while human TRIM5alpha has a short half-life and turns over quickly in cells, TRIM34 is significantly more stable (Li et al., 2007). It may be that TRIM34 stabilizes TRIM5alpha, thereby allowing it to restrict HIV capsids that transit differently than wild type into the nucleus. In this model, wild type HIV-1 capsids are not sensitive to restriction as they transit the cytoplasm more efficiently and/or via a different nuclear entry route that allows escape from restriction. As proposed by Sultana et al., an accelerated rate of uncoating could be a mechanism of escape from cellular restrictions, particularly those that target the HIV-1 capsid (Sultana et al., 2019).

Either of these models are sufficient to explain why TRIM34 is required for restriction of the N74D virus and why this restriction depends on the presence of TRIM5alpha. Work is underway to test these models and provide insight into the mechanism of TRIM34 restriction.

## Acknowledgments

We thank Masa Yamashita for providing capsid mutant proviral plasmids, including the unpublished P90A provirus, Daryl Humes for help with CD4 T cell preparations and electroporations, Sydney Fine for performing editing analysis on edited T cell pools and the Fred Hutch Genomics Core, including Jeff Delrow and Ryan Basom, for help with deep sequencing and HIV-CRISPR screen analysis. We are also grateful to the blood donors who contributed cells to this study. This work was supported by NIH grants: R01 AI147877 (M.O.), CFAR AI027757 (M.O.), R01 AI30927 (M.E), 1R37AI147868 (J.L.) and R01 AI093258 (E.C.).

## References

Achuthan, V., Perreira, J.M., Sowd, G.A., Puray-Chavez, M., McDougall, W.M., Paulucci-Holthauzen, A., Wu, X., Fadel, H.J., Poeschla, E.M., Multani, A.S., et al. (2018). Capsid-CPSF6 Interaction Licenses Nuclear HIV-1 Trafficking to Sites of Viral DNA Integration. Cell Host Microbe 24, 392–404 e398.

Ambrose, Z., and Aiken, C. (2014). HIV-1 uncoating: connection to nuclear entry and regulation by host proteins. Virology 454-455, 371–379.

Ambrose, Z., Lee, K., Ndjomou, J., Xu, H., Oztop, I., Matous, J., Takemura, T., Unutmaz, D., Engelman, A., Hughes, S.H., et al. (2012). Human immunodeficiency virus type 1 capsid mutation N74D alters cyclophilin A dependence and impairs macrophage infection. J Virol 86, 4708–4714.

Asmuth, D.M., Murphy, R.L., Rosenkranz, S.L., Lertora, J.J., Kottilil, S., Cramer, Y., Chan, E.S., Schooley, R.T., Rinaldo, C.R., Thielman, N., et al. (2010). Safety, tolerability, and mechanisms of antiretroviral activity of pegylated interferon Alfa-2a in HIV-1-monoinfected participants: a phase II clinical trial. J Infect Dis 201, 1686–1696.

Bejarano, D.A., Peng, K., Laketa, V., Borner, K., Jost, K.L., Lucic, B., Glass, B., Lusic, M., Muller, B., and Krausslich, H.G. (2019). HIV-1 nuclear import in macrophages is regulated by CPSF6-capsid interactions at the nuclear pore complex. Elife 8.

Bulli, L., Apolonia, L., Kutzner, J., Pollpeter, D., Goujon, C., Herold, N., Schwarz, S.M., Giernat, Y., Keppler, O.T., Malim, M.H., et al. (2016). Complex Interplay between HIV-1 Capsid and MX2-Independent Alpha Interferon-Induced Antiviral Factors. J Virol 90, 7469–7480.

Campbell, E.M., and Hope, T.J. (2015). HIV-1 capsid: the multifaceted key player in HIV-1 infection. Nat Rev Microbiol 13, 471–483.

De Iaco, A., and Luban, J. (2011). Inhibition of HIV-1 infection by TNPO3 depletion is determined by capsid and detectable after viral cDNA enters the nucleus. Retrovirology 8, 98.

Duggal, N.K., and Emerman, M. (2012). Evolutionary conflicts between viruses and restriction factors shape immunity. Nat Rev Immunol 12, 687–695.

Finzi, D., Blankson, J., Siliciano, J.D., Margolick, J.B., Chadwick, K., Pierson, T., Smith, K., Lisziewicz, J., Lori, F., Flexner, C., et al. (1999). Latent infection of CD4+ T cells provides a mechanism for lifelong persistence of HIV-1, even in patients on effective combination therapy. Nat Med 5, 512–517.

Ganser-Pornillos, B.K., and Pornillos, O. (2019). Restriction of HIV-1 and other retroviruses by TRIM5. Nat Rev Microbiol.

Hatziioannou, T., Princiotta, M., Piatak, M., Jr., Yuan, F., Zhang, F., Lifson, J.D., and Bieniasz, P.D. (2006). Generation of simian-tropic HIV-1 by restriction factor evasion. Science 314, 95.

Henning, M.S., Dubose, B.N., Burse, M.J., Aiken, C., and Yamashita, M. (2014). In vivo functions of CPSF6 for HIV-1 as revealed by HIV-1 capsid evolution in HLA-B27-positive subjects. PLoS Pathog 10, e1003868.

Hilditch, L., and Towers, G.J. (2014). A model for cofactor use during HIV-1 reverse transcription and nuclear entry. Curr Opin Virol 4, 32–36.

Hrecka, K., Hao, C., Gierszewska, M., Swanson, S.K., Kesik-Brodacka, M., Srivastava, S., Florens, L., Washburn, M.P., and Skowronski, J. (2011). Vpx relieves inhibition of HIV-1 infection of macrophages mediated by the SAMHD1 protein. Nature 474, 658–661.

Hulme, A.E., Kelley, Z., Okocha, E.A., and Hope, T.J. (2015). Identification of capsid mutations that alter the rate of HIV-1 uncoating in infected cells. J Virol 89, 643–651.

Jimenez-Guardeno, J.M., Apolonia, L., Betancor, G., and Malim, M.H. (2019). Immunoproteasome activation enables human TRIM5alpha restriction of HIV-1. Nat Microbiol 4, 933–940.

Kim, K., Dauphin, A., Komurlu, S., McCauley, S.M., Yurkovetskiy, L., Carbone, C., Diehl, W.E., Strambio-De-Castillia, C., Campbell, E.M., and Luban, J. (2019). Cyclophilin A protects HIV-1 from restriction by human TRIM5alpha. Nat Microbiol.

Laguette, N., Sobhian, B., Casartelli, N., Ringeard, M., Chable-Bessia, C., Segeral, E., Yatim, A., Emiliani, S., Schwartz, O., and Benkirane, M. (2011). SAMHD1 is the dendritic- and myeloid-cell-specific HIV-1 restriction factor counteracted by Vpx. Nature 474, 654–657.

Lee, K., Ambrose, Z., Martin, T.D., Oztop, I., Mulky, A., Julias, J.G., Vandegraaff, N., Baumann, J.G., Wang, R., Yuen, W., et al. (2010). Flexible use of nuclear import pathways by HIV-1. Cell Host Microbe 7, 221–233.

Li, X., Gold, B., O’HUigin, C., Diaz-Griffero, F., Song, B., Si, Z., Li, Y., Yuan, W., Stremlau, M., Mische, C., et al. (2007). Unique features of TRIM5alpha among closely related human TRIM family members. Virology 360, 419–433.

Li, X., Yeung, D.F., Fiegen, A.M., and Sodroski, J. (2011). Determinants of the higher order association of the restriction factor TRIM5alpha and other tripartite motif (TRIM) proteins. J Biol Chem 286, 27959–27970.

Luban, J., Bossolt, K.L., Franke, E.K., Kalpana, G.V., and Goff, S.P. (1993). Human immunodeficiency virus type 1 Gag protein binds to cyclophilins A and B. Cell 73, 1067–1078.

Malim, M.H., and Bieniasz, P.D. (2012). HIV Restriction Factors and Mechanisms of Evasion. Cold Spring Harb Perspect Med 2, a006940.

Mariani, R., Chen, D., Schrofelbauer, B., Navarro, F., Konig, R., Bollman, B., Munk, C., Nymark-McMahon, H., and Landau, N.R. (2003). Species-specific exclusion of APOBEC3G from HIV-1 virions by Vif. Cell 114, 21–31.

McCauley, S.M., Kim, K., Nowosielska, A., Dauphin, A., Yurkovetskiy, L., Diehl, W.E., and Luban, J. (2018). Intron-containing RNA from the HIV-1 provirus activates type I interferon and inflammatory cytokines. Nat Commun 9, 5305.

Nisole, S., Stoye, J.P., and Saib, A. (2005). TRIM family proteins: retroviral restriction and antiviral defence. Nat Rev Microbiol 3, 799–808.

OhAinle, M., Helms, L., Vermeire, J., Roesch, F., Humes, D., Basom, R., Delrow, J.J., Overbaugh, J., and Emerman, M. (2018). A virus-packageable CRISPR screen identifies host factors mediating interferon inhibition of HIV. Elife 7.

Orimo, A., Tominaga, N., Yoshimura, K., Yamauchi, Y., Nomura, M., Sato, M., Nogi, Y., Suzuki, M., Suzuki, H., Ikeda, K., et al. (2000). Molecular cloning of ring finger protein 21 (RNF21)/interferon-responsive finger protein (ifp1), which possesses two RING-B box-coiled coil domains in tandem. Genomics 69, 143–149.

Pertel, T., Hausmann, S., Morger, D., Zuger, S., Guerra, J., Lascano, J., Reinhard, C., Santoni, F.A., Uchil, P.D., Chatel, L., et al. (2011). TRIM5 is an innate immune sensor for the retrovirus capsid lattice. Nature 472, 361–365.

Price, A.J., Fletcher, A.J., Schaller, T., Elliott, T., Lee, K., KewalRamani, V.N., Chin, J.W., Towers, G.J., and James, L.C. (2012). CPSF6 defines a conserved capsid interface that modulates HIV-1 replication. PLoS Pathog 8, e1002896.

Rasaiyaah, J., Tan, C.P., Fletcher, A.J., Price, A.J., Blondeau, C., Hilditch, L., Jacques, D.A., Selwood, D.L., James, L.C., Noursadeghi, M., et al. (2013). HIV-1 evades innate immune recognition through specific cofactor recruitment. Nature 503, 402–405.

Rasheedi, S., Shun, M.C., Serrao, E., Sowd, G.A., Qian, J., Hao, C., Dasgupta, T., Engelman, A.N., and Skowronski, J. (2016). The Cleavage and Polyadenylation Specificity Factor 6 (CPSF6) Subunit of the Capsid-recruited Pre-messenger RNA Cleavage Factor I (CFIm) Complex Mediates HIV-1 Integration into Genes. J Biol Chem 291, 11809–11819.

Saito, A., Henning, M.S., Serrao, E., Dubose, B.N., Teng, S., Huang, J., Li, X., Saito, N., Roy, S.P., Siddiqui, M.A., et al. (2016). Capsid-CPSF6 Interaction Is Dispensable for HIV-1 Replication in Primary Cells but Is Selected during Virus Passage In Vivo. J Virol 90, 6918–6935.

Sawyer, S.L., Emerman, M., and Malik, H.S. (2007). Discordant evolution of the adjacent antiretroviral genes TRIM22 and TRIM5 in mammals. PLoS Pathog 3, e197.

Sawyer, S.L., Wu, L.I., Emerman, M., and Malik, H.S. (2005). Positive selection of primate TRIM5alpha identifies a critical species-specific retroviral restriction domain. Proc Natl Acad Sci U S A 102, 2832–2837.

Schaller, T., Ocwieja, K.E., Rasaiyaah, J., Price, A.J., Brady, T.L., Roth, S.L., Hue, S., Fletcher, A.J., Lee, K., KewalRamani, V.N., et al. (2011). HIV-1 capsid-cyclophilin interactions determine nuclear import pathway, integration targeting and replication efficiency. PLoS Pathog 7, e1002439.

Shah, V.B., Shi, J., Hout, D.R., Oztop, I., Krishnan, L., Ahn, J., Shotwell, M.S., Engelman, A., and Aiken, C. (2013). The host proteins transportin SR2/TNPO3 and cyclophilin A exert opposing effects on HIV-1 uncoating. J Virol 87, 422–432.

Sowd, G.A., Serrao, E., Wang, H., Wang, W., Fadel, H.J., Poeschla, E.M., and Engelman, A.N. (2016). A critical role for alternative polyadenylation factor CPSF6 in targeting HIV-1 integration to transcriptionally active chromatin. Proc Natl Acad Sci U S A 113, E1054–1063.

Sultana, T., Mamede, J.I., Saito, A., Ode, H., Nohata, K., Cohen, R., Nakayama, E.E., Iwatani, Y., Yamashita, M., Hope, T.J., et al. (2019). Multiple pathways to avoid IFN-beta sensitivity of HIV-1 by mutations in capsid. J Virol.

Yamashita, M., and Emerman, M. (2004). Capsid is a dominant determinant of retrovirus infectivity in nondividing cells. J Virol 78, 5670–5678.

Yamashita, M., and Engelman, A.N. (2017). Capsid-Dependent Host Factors in HIV-1 Infection. Trends Microbiol.

Yang, Y., Brandariz-Nunez, A., Fricke, T., Ivanov, D.N., Sarnak, Z., and Diaz-Griffero, F. (2014). Binding of the rhesus TRIM5alpha PRYSPRY domain to capsid is necessary but not sufficient for HIV-1 restriction. Virology 448, 217–228.

Zhang, F., Hatziioannou, T., Perez-Caballero, D., Derse, D., and Bieniasz, P.D. (2006). Antiretroviral potential of human tripartite motif-5 and related proteins. Virology 353, 396–409.

